# Integrated biphasic growth rate, gene expression, and cell-size homeostasis behaviour of single *B. subtilis* cells

**DOI:** 10.1101/510925

**Authors:** Niclas Nordholt, Johan H. van Heerden, Frank J. Bruggeman

## Abstract

The growth rate of single bacterial cells is continuously disturbed by random fluctuations in biosynthesis rates and by deterministic cell-cycle events, such as division, genome duplication, and septum formation. It is not understood whether, and how, bacteria reject these disturbances. Here we quantified growth and constitutive protein expression dynamics of single *Bacillus subtilis* cells, as a function of cell-cycle-progression. Variation in birth size and growth rate, resulting from unequal cell division, is largely compensated for when cells divide again. We analysed the cell-cycle-dynamics of these compensations and found that both growth and protein expression exhibited biphasic behaviour. During a first phase of variable duration, the absolute rates were approximately constant and cells behaved as sizers. In the second phase, rates increased and growth behaviour exhibited characteristics of a timer-strategy. This work shows how cell-cycle-dependent rate adjustments of biosynthesis and growth are integrated to compensate for physiological disturbances caused by cell division.

**IMPORTANCE:** Under constant conditions, bacterial populations can maintain a fixed average cell size and constant exponential growth rate. At the single cell-level, however, cell-division can cause significant physiological perturbations, requiring compensatory mechanisms to restore the growth-related characteristics of individual cells toward that of the average cell. Currently, there is still a major gap in our understanding of the dynamics of these mechanisms, i.e. how adjustments in growth, metabolism and biosynthesis are integrated during the bacterial cell-cycle to compensate the disturbances caused by cell division. Here we quantify growth and constitutive protein expression in individual bacterial cells at sub-cell-cycle resolution. Significantly, both growth and protein production rates display structured and coordinated cell-cycle-dependent dynamics. These patterns reveal the dynamics of growth rate and size compensations during cell-cycle progression. Our findings provide a dynamic cell-cycle perspective that offers novel avenues for the interpretation of physiological processes that underlie cellular homeostasis in bacteria.

## Introduction

Under constant conditions, isogenic populations of bacteria maintain time-invariant distributions of cell size, generation time and macromolecular composition. This growth mode is called balanced growth^1,2,3,4^. Yet, individual cells display molecular fluctuations that are the source for non-genetic heterogeneity within an isogenic population^5,6,7,8,9^ and influence single cell growth behaviour^10^. How bacteria achieve the required homeostasis associated with balanced growth has been the subject of decades of microbial studies, both at the level of populations^11^ and single-cells^12,13,14,15,16,17^. Nowadays, thousands of individual cells can be studied simultaneously, at high temporal resolution using fluorescence microscopy, which led to the discovery of principles of cell-size homeostasis^13,14,15,16,17^, and opens up new opportunities for more quantitative analysis of the dynamics of a single cell along its cell-cycle.

One explanation of steady-state exponential growth of a population of bacterial cells is that its constituent single cells grow at the same constant exponential rate that fluctuates^10^ independent of the cell-cycle. However, it might also be different. The physiological state of a cell might be continuously perturbed to give rise to deviations from exponential growth that are similar across cells; for instance, because they are cell-cycle stage dependent. Since single-cell growth is asynchronous in a population of cells, those deviations would not be evident from population studies. Therefore, studies on growth rate of single cells are required.

Cell-cycle dependent phenomena that can in principle cause systematic growth rate perturbations have been described. Genome replication can influence the expression of biosynthetic genes by increasing gene copy numbers, which can cause changes in biosynthesis and growth rate^18^. Since the volume of a cell increases faster than its surface area^19^, the demands for membrane components changes with cell-cycle progression. The growth of the cell wall occurs continuously and uniformly along the cell axis in rod-shaped bacteria^20^, but the rate of peptidoglycan (PG) incorp oration accelerates at the constriction site, which leads to an increased demand for PG precursors^21,22^. Thus, the demands for particular biosynthetic precursors are dependent on cell-cycle progression, which requires adjustments of the metabolic activity of the cell.

Cell-cycle dependent changes in cell structure, metabolic activity and protein expression beg the question whether the growth rate of an individual cell is at all times proportional to its size, and therefore exponential. Or is it rather adjusted continuously in response to cell-cycle dependent perturbations? Do distinct growth phases occur, associated with passing of checkpoints and completion of cell-cycle-specific events? Early observations were made that suggest the existence of such distinct growth phases^12,23,24^. However, these studies relied on the inference of single-cell growth rates from stationary cell-size distributions. Few contemporary attempts have been made to characterise bacterial growth at a sub-cell-cycle resolution^25,26^, with divergent conclusions relying on quantifications of only a small number of cells.

Here we characterise the elongation rate dynamics of thousands of individual B. *subtilis* cells with sub-cell-cycle resolution, using time-lapse fluorescence microscopy, under three growth conditions with different population growth rates, which we varied through the addition of different carbon sources. Cell-cycle progression in *B. subtilis* is characterised by two distinct growth phases. A first phase, during which cells behave as sizers and grow at a constant absolute elongation rate. And a second phase, during which cells act as timers and the absolute elongation rate increases. The specific-growth-rate dynamics of a single cell depends on its size at birth. Across conditions, cells that are born smaller than average initially have a higher specific elongation rate than those that are born larger. At the end of the cell-cycle, cells display cell-size independent growth rates and cell-size disturbances have largely been compensated for. We find that cells maintain stable protein concentrations due to synchronous adjustments in protein production and dilution (i.e. growth) rates.

## Results

### Non-exponential, single-cell growth: cell division generates a size-dependent specific elongation rate

Exponential growth of cell volume by rod-shaped cells, such as *B. subtilis*, at a fixed rate corresponds to a non-linear relation of the specific elongation rate (in terms of cell length) with cell-cycle progression (see Appendix). Since we quantify cell growth in terms of cell length, we have to consider deviations from this non-linear relation to identify deviations from exponential growth. We distinguish two growth rates of the length (*L*) of the cell: the (absolute) elongation rate of the cell 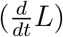 and the specific elongation rate of the cell 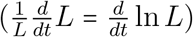. We focus on: i. the specific elongation rate of a single cell because a cell population that grows balanced has a constant specific growth rate and on ii. the absolute elongation rate because this is proportional to the current metabolic activity of the cell. The experimental data in this paper are all valid at balanced growth of the population of cells; the probability distributions of various cell characteristics were tested for their time invariance (Figs. S6-S7). In addition, we evaluated whether observed growth patterns are robust to measurement noise by comparing measured and simulated single-cell growth profiles (see Appendix, Figs. S1-S5).

Figure 1A shows the average growth behaviour of the length of single B. *subtilis* cells at three different conditions, measured on agarose pads, using real-time imaging of cell growth. We observe systematic deviations from exponential growth that are dependent on cell-cycle progression, cell size and nutrient conditions (Fig. 1B,C). The specific elongation rate 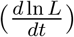 displays large systematic deviations from its expected value (the dotted lines in Fig. 1B) of up to ±20% at the start, halfway and the end of the cell-cycle. Thus, even though the population grows with a time-invariant specific elongation rate, individual cells do not maintain this value along their cell-cycle. They experience perturbations from the metabolic steady state that is required for a constant specific elongation rate. Those perturbations are somehow associated with cell-cycle progression and occur in all cells similarly; otherwise, one would not observe systematic deviations from exponential growth when the single-cell data is averaged, as in Fig. 1.

**Figure 1:**
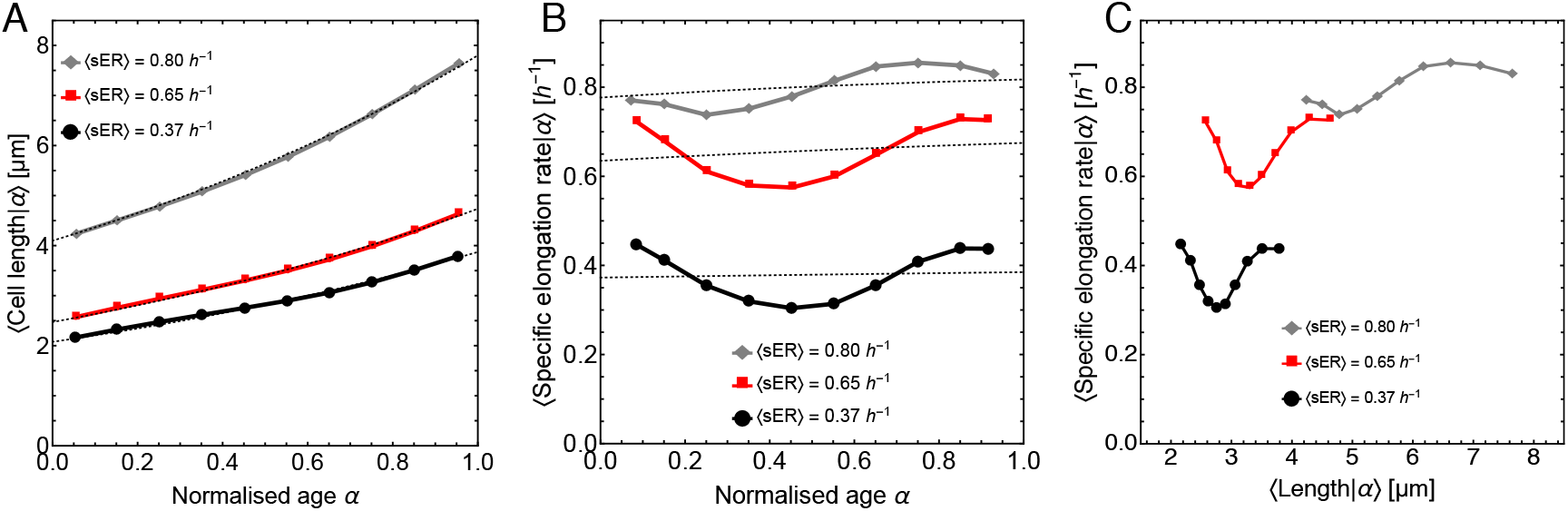
Single cells show systematic deviations from exponential growth along their cell-cycles. **A.** The average length of a single cell as function of cell-cycle progression is shown. Cell-cycle progression can be expressed in terms of normalised cell age (*α*), which is defined as the time elapsed since birth divided by the generation time of a cell, i.e. it equals 0 at birth and 1 at division. The cell growth data approximates the expected theoretical relation (dotted lines; see Appendix). **B.** The average specific elongation rate (*sER*) of a single cell as function of its normalised age. If growth is exponential, sER follows the relation shown in the Appendix (dotted lines). **C.** The average specific elongation rate (*sER*) of a single cell as function of its length. Panels A to C consider only those cells for which we observed both their birth and division event. Cell length or specific elongation rate was conditioned on normalised cell age into bins of width 0.1 and the mean value for each bin was calculated. We remark that the standard-error bars are smaller than the plot markers. In total, we studied *N* = 15891 cells at the slow growth condition (0.37 *hr*^−1^, arabinose), 12553 cells at an intermediate growth rate (0.65 *hr*^−1^, glucose) and 2887 cells at fast growth rate (0.80 *hr*^−1^, glucose + 4 amino acids condition: methionine, histidine, glutamate and tryptophan). Cells were pooled from 5 independent experiments per condition (see Methods). Distributions of birth length and division length were time-invariant over several generations, confirming balanced growth of the cell population (Fig. S7). We note that the specific growth rate averaged over all cells (〈*sER*〉) was in excellent agreement with the respective population growth rate, for each condition (Fig. S6).

To better understand the origins of the systematic deviations from exponential growth, we partitioned the cell data into 5 bins according to their birth size (Fig. 2A,B). Each class contains 20% of all the studied cells. We considered only those cells for which complete cell-cycles (birth and division) were observed. Figure 2A shows the frequency distributions of birth and division length, coloured according to the different classes, for the intermediate growth rate condition (0.65 *hr*^−1^, glucose). The birth length distributions are separated, but the division length distributions overlap, indicating that a compensating mechanism for cell size homeostasis exists. Cells that are born larger or smaller than average, have, on average, shorter and longer generation times (Fig. 2A, distributions on x-axis), respectively.

**Figure 2:**
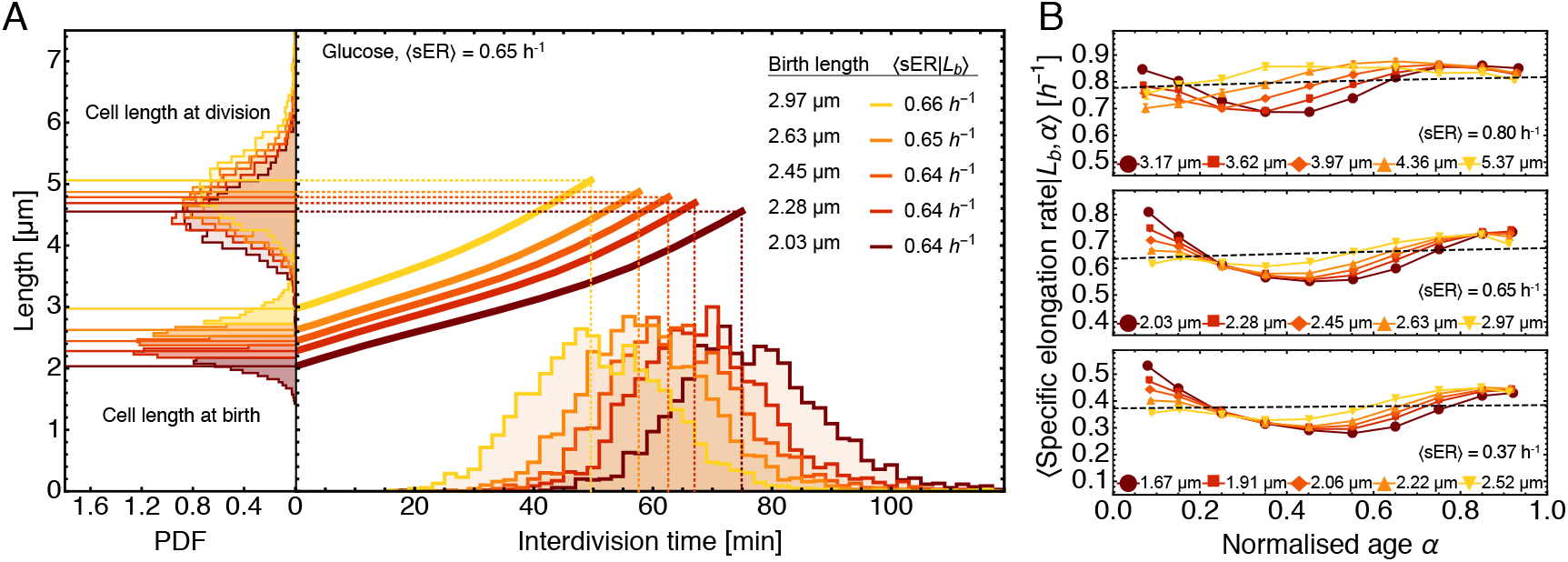
Cell division causes size and elongation rate disturbances from average cell behaviour. **A.** Average, single-cell length is shown as function of time, classified according to five birth length classes, each containing 20% of all the studied cells. Data is shown for the intermediate growth rate condition (0.65 *hr*^−1^, glucose). The frequency distributions along the y-axis show the length distributions at birth (left bottom) and division (left top). Generation time distributions of the individual birth length classes are shown on the x-axis. Horizontal and vertical dashed lines indicate the mean division length and interdivision time of each birth length class, respectively. **B.** Specific elongation rate *sER* of different birth length classes as function of cell age for fast (0.80 *hr*^−1^, top), intermediate (0.65 *hr*^−1^, middle) and slow (0.37 *hr*^−1^, bottom) growth. The legends indicate the mean birth length of the respective birth length class. The dashed line is the specific elongation rate according to equation 2 and the same as in Fig. 1A. Note that the mean 〈*sER*|*L_b_, α*〉 for the different birth length classes was the same as the population average, 〈*sER*〉. The number of cells per birth length class: *N* = 3179, 2511, 578 for slow, intermediate and fast growth, respectively.

The specific elongation rates averaged across the entire cell-cycle hardly differs between the cell classes (inset table in Fig. 2A). This picture changes at the sub-cell-cycle resolution. We determined how the average specific elongation rates within the cell classes varied along the cell-cycle. Figure 2B indicates that the cells of each birth-size class display systematic deviations from exponential growth at the start, halfway and the end of the cell-cycle. This figure indicates that cell division perturbs the specific elongation rate in a birth-size dependent manner. Cells that are born comparatively small, generally deviate most from exponential growth (Fig. 2B) and initially have a higher specific elongation rate than larger cells. Figure 2B also shows that although significant deviations occur in elongation rate at birth, all cells converge to nearly the same specific elongation rate at the end of the cell-cycle (*α* ≈ 1), regardless of their cell size at birth and their starting elongation rate, and that growth rate heterogeneity has decreased (Fig. S8).

Thus, it appears that cell division disturbs the metabolic steady-state in a daughter cell such that its elongation rate deviates from its mother, and smaller cells grow faster than larger cells. The specific elongation rate exhibits birth-size dependent dynamic changes during the cell-cycle until, at cell division, all cells have a birth-size independent growth rate. Cell division, therefore, leads to a cell-size dependent perturbation of the specific elongation rate of a cell that is compensated for during cell-cycle progression.

### Non-exponential single-cell growth: absolute elongation rate is biphasic with cell-cycle progression

While figure 2 analysed the behaviour of the specific elongation rate 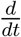 ln *L*, figure 3 focusses on the absolute elongation rate 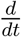*L*. Its dynamics along the cell-cycle displays biphasic behaviour (Fig. 3). When plotted as function of cell-cycle progression, two phases appear to exist. The first phase is characterised by an approximately constant absolute elongation rate (implying linear growth). The second phase is characterised by an increasing absolute elongation rate (as expected for exponential growth). We note that the second phase occurs later in the cell-cycle (i.e. the first phase is longer) in cells that were small at birth (Fig. 3A). An exception to the biphasic growth pattern is observed only for cells that were born larger than average in the fast growth condition (top panel in Fig. 3A, two largest size bins: 5.37 and 4.36 *μ*m). The elongation rate of these cells increases throughout the entire cell-cycle.

**Figure 3:**
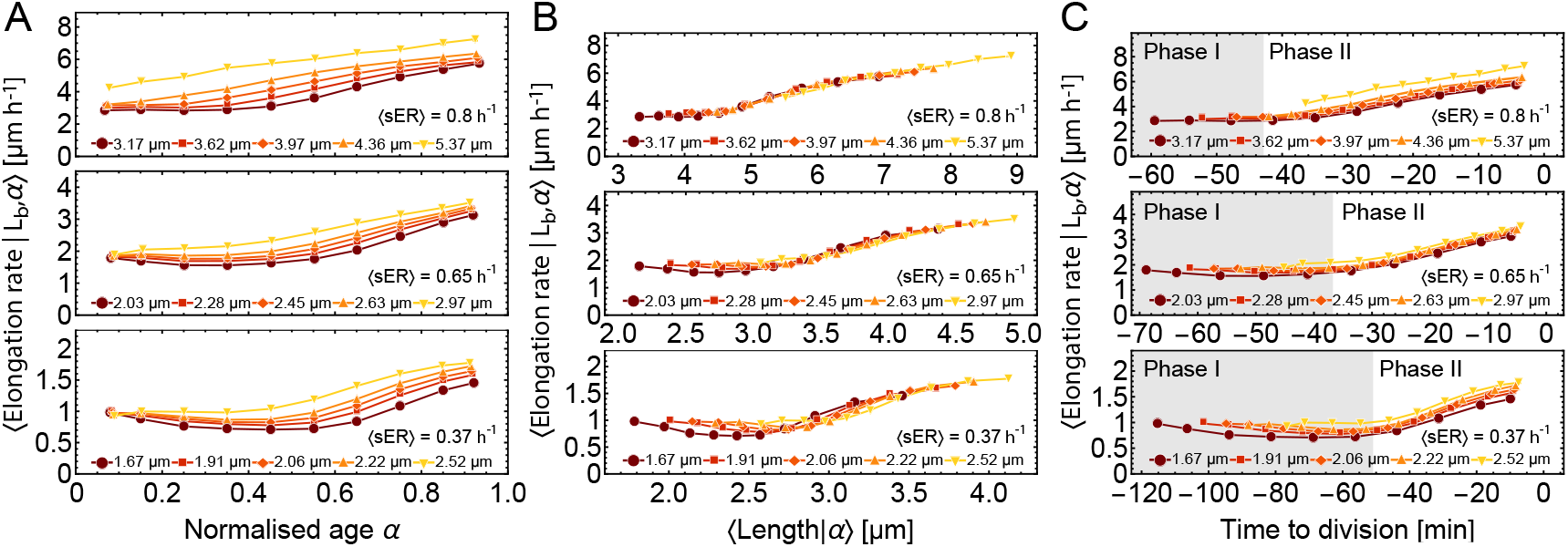
Single-cell growth as function of the cell-cycle is biphasic. **A.** The absolute elongation rate as function of the normalised cell age indicates that most cells start with a fairly constant elongation rate that increases as time progresses in a birth-size dependent manner. **B.** The absolute elongation rate depends on cell length, which is weakly birth-size dependent when cells grew in the slow and intermediate growth condition. **C.** The absolute elongation rate of single cells displayed as function of the time to division reveals that all cells, within a specific condition, start growing faster at an approximately fixed time before they divide, indicating biphasic growth, with phase I marked in grey. Below we analyse this data more carefully to confirm that growth is biphasic. At the fast growth condition, upper plot, the cells that were born large appear to skip phase I.

To understand whether the differences in the duration of the two phases are related to cell length, we plot the elongation rate as function of cell length (Fig. 3B). These curves deviate from the expected exponential curve and the effect of birth size is greatest at the slow-growth condition. The overall curvature is very similar across the three growth conditions: it is biphasic, first the elongation rate is fairly constant while it rises in the second phase.

The generation time of cells correlates strongly with their size at birth (Fig. S9). We therefore tested whether the onsets between the first and second phases of growth for different birth-size classes (Fig 3A,B) relate to absolute differences in cell-cycle duration. It seems reasonable to expect that any homeostatic adjustments made during the cell-cycle function to ensure compensation of physiological disturbances (e.g. cell size differences) generated at birth. This makes cell division a logical, albeit non-conventional, anchoring point for evaluation of cell-cycle-related growth or gene expression dynamics. We evaluated the average elongation rate of cells in the five different birth-size classes as function of the time-to-division (rather than time-since-birth). Interestingly, from this perspective we found alignment of phase transitions for different birth size-classes, with rates increasing at a constant condition-dependent time before cell division, independent of birth length (Fig. 3C). By fitting a piecewise function to the data (see Methods), we estimated the point of phase transition at slow, intermediate and fast growth to be 51.3±1.2, 37.1±1 and 43.4±0.7 minutes (Means of the birth size classes ± S.D.) prior to division, respectively (Fig. 3C). Below we analyse the same data in a different manner and conclude the existence of two phases from the perspective of mechanisms of cell-size homeostasis.

We conclude that the elongation rate of cells as function of the cell-cycle is not exponential, it appears to be biphasic and birth-size dependent. For a specific growth condition, cells start the second phase of growth at a similar time before the cell division, regardless of their birth size.

### Strong correlation between specific protein synthesis rate and specific elongation growth

The observed biphasic, non-exponential growth behaviour of a single cell along its cell-cycle raises the question of how its protein synthesis rate behaves. To address this, we exploited the fact that the *B. subtilis* strain used in this study expressed a constitutive fluorescent protein (GFP), from a genomic locus, that is controllable by an inducible promoter; we described this strain previously^8^. In this section we analyse the strain grown at full induction with 1 mM IPTG. This dataset constitutes a subset of the data shown in figures 1 - 3 (see Methods).

In Fig. 4A, the average elongation rate and the average fluorescence production rate are both plotted as function of the time to division. Both rates correlate strongly across the two phases. This is perhaps not so surprising, as volume growth rate is often viewed as a result of the synthesis of proteins that occupy volume^27^. Figure 4B indicates that the correlation between the elongation and fluorescence production rate is preserved in the birth-size classes. Figure 4C displays the strong correlation of the elongation rate and the fluorescence production rate (both normalised with respect to the mean value across all conditions).

**Figure 4:**
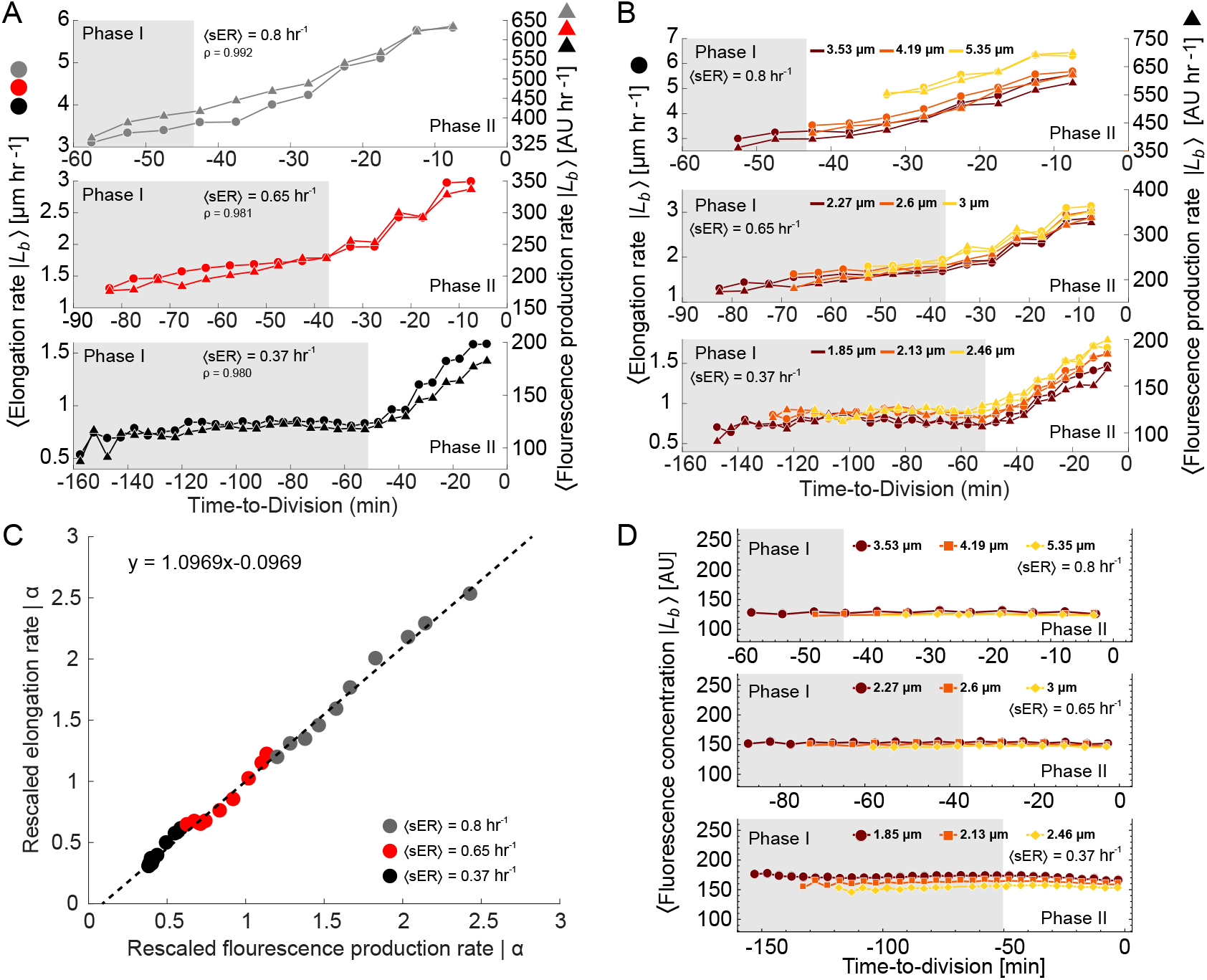
The fluorescence production rate of single cells as function of the cell-cycle correlates strongly with their elongation rate. **A.** Average absolute elongation rate and fluorescence production rate as function of the time to division. Phase I is indicated in grey. The three growth conditions are shown on top of each other, with the lowest growth rate condition at the bottom. The elongation rate and the fluorescence production rate of cells show near identical behaviour. The Spearman correlation coefficient *ρ* between the average fluorescence production rate and the average elongation rate is indicated in the plots. Number of cells that express GFP N= 4233, 1778, 577 for slow, intermediate and fast growth, respectively. **B.** Average absolute elongation rate and fluorescence production rate as function of the time to division for five cell classes binned according to their birth size ranges. The average growth rate and the average fluorescence production rate of cells show near identical behaviour, regardless of their birth size. Each birth size class contains one third of all GFP expressing cells from each condition. **C.** The elongation rate as function of the fluorescence production rate for all experiments, normalised with respect to their mean values across all experiments, indicates a near linear relationship. **D.** The average fluorescence concentration in cells is independent of the time to division and, therefore, also of cell-cycle progression. For plots A, B, D cells were conditioned on time-to-division into bins of 5 minute width and the average value of each bin with > 50 observations was computed.

Since the protein production rate and the elongation rate, responsible for protein dilution, strongly correlate one would expect that the protein concentration in cells remains constant along the cell-cycle. Figure 4D confirms this expectation. We note that this expectation is only valid for stable proteins, which are not rapidly degraded, like the fluorescent protein we used.

Thus, we can conclude that cells maintain protein-concentration homeostasis (for stable proteins) along their cell-cycles, because the balance between protein synthesis and degradation is unaffected by growth-rate changes; since the synthesis and degradation rate (i.e. the dilution rate) strongly correlate across the cell-cycle of a cell. This likely results from protein synthesis being the main determinant of the cellular growth rate. These findings also suggests that all cell-cycle dependent changes in growth rate are preceded by changes in protein synthesis rate, which is in agreement with Kiviet & Tans^10^.

### Cells initially grow as sizers after birth, then become timers until they divide, behaving overall as imperfect adders

To address how *B. subtilis* achieves size homeostasis along its cell-cycle, we determined whether it behaves as a sizer, timer or adder in each cell-cycle phases and what the overall behaviour is. Figure 2A indicates that the cell size added during an entire cell-cycle correlates with cell size at birth with small-born cells adding more than large-born cells. This is confirmed when we determine B. *subtilis*’ cell size homeostasis mechanism (Fig. 5A) during its entire cell-cycle, which exhibits mixed sizer-adder behaviour; an imperfect adder. Figure 5B and C indicate that *B. subtilis* ‘ behaviour in the first cell-cycle phase resembles that of a sizer. Figure 5B shows that the coefficient of variation in cell length decreases immediately after cell birth until, for the slow and intermediate growth conditions, it reaches a minimum, which coincides with the onset of phase II as we determined with the piecewise function method (described in the Methods). Thus, during phase I, the cells act as sizers; since the variation in length evident at cell birth is reduced during this period. This is also evident from Fig. 5C, which indicates that cells add, in phase I, an amount of length that compensates for their length difference with respect to the mean cell.

**Figure 5:**
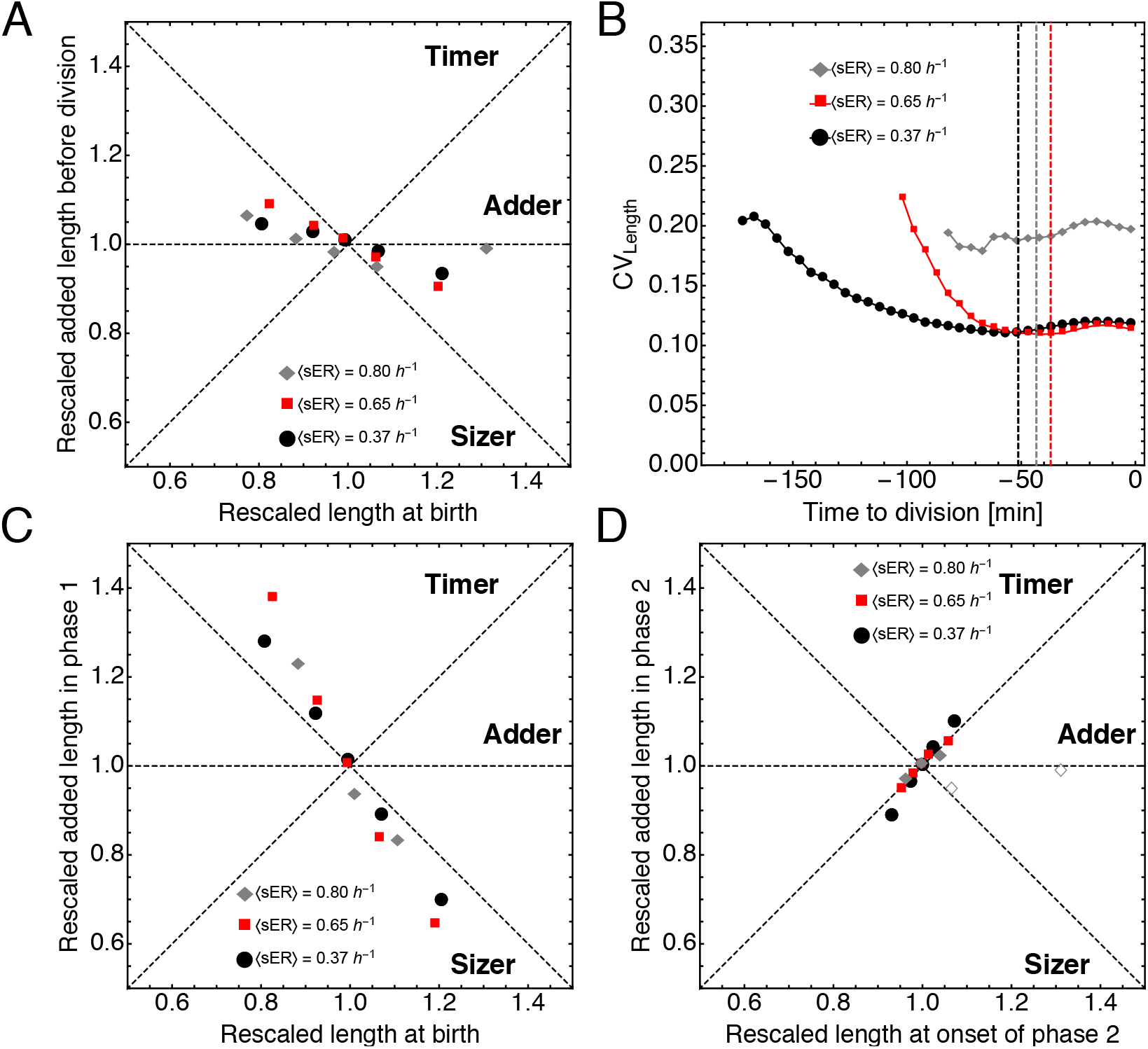
Adder behaviour is a consequence of a sizer in phase I and a subsequent timer. **A.** Correlations between rescaled cell size at birth and rescaled added length before division identify *B. subtilis* as an imperfect adder over its cell cylce^28^. Each symbol represents one of the five birth-size classes that were defined in Fig. 2. **B.** Coefficient of variation in cell length plotted as function of time to division. The coefficient of variation decreases immediately after birth and reaches a minimum approximately at the end of phase I (dashed lines), after which it increases again. **C.** Cells behave as sizers until the end of phase I. The two largest bins from the fast growth condition are excluded as they lack phase I as determined from Fig. 3C. **D.** At the onset of phase II, cells grow for an approximately constant amount of time until they divide, resulting in timer behaviour. The two largest bins from the fast growth conditions are displayed as empty symbols and are the same as in A.

Another conclusion from Fig. 5B is that cell length heterogeneity is largely generated by cell division, as the coefficient of variation in cell length is largest at birth, and much greater than its value at division. Finally, we studied the cell-size homeostasis mechanism in the second phase of the cell-cycle (Fig. 5C), which is timer like, indicating that the duration of phase II is independent of the cell length at the start of phase II; as would be the case if some cellular process with a fairly fixed duration is initiated, and completed. This process might by DNA replication, septum formation and cell division, because at the high growth rate condition it is expected that most cells commence with DNA replication soon or immediately after cell division, resulting in a short or even absent phase I. At the high growth-rate condition, we did indeed see that the largest and fastest growing cells lack a phase I and behave as adders during their cell-cycle.

Thus, depending on the growth condition *B. subtilis* cells grow biphasically, as sizers in the first phase and as timers in the second, and overall as imperfect adders (with a weak sizer component). Cells that are born small compensate for their length deficit in the first phase, while large newborn cells grow less than average. The duration of the second phase is comparable across cells, except for those cells that were born larger than the size threshold for the start of phase II, in the fast growth condition.

## Discussion

Our experiments indicate that an isogenic population of *B. subtilis* cells that grows at a constant specific growth rate consists of individual cells that show cell-cycle dependent deviations from this fixed rate (Fig. 1). This behaviour was observed at three conditions. These systematic growth-rate deviations are not evident at the population level because individual cells pass asynchronously through their cell-cycles. Figure 2B illustrates four important effects: i. growth rate variation is introduced at cell birth (see also Fig. S8), ii. smaller cells grow faster at birth, iii. the specific growth rate dynamics of a single cell depends on its birth size, and iv. all cells finally reach a specific growth rate that is independent of their birth size at the end of the cell-cycle. Cell size heterogeneity is also greatly influenced by cell division (Fig. 5B), its coefficient of variation is large at cell birth and the size variation is compensated for during the cell-cycle, which is indicated in Fig. 2A and in Fig. 5A, where it can be seen that the total added length during a cell-cycle corrects for variations in birth size.

The dynamic compensation of growth rate and size variation during the cell-cycle might be the outcome of a regulatory mechanism that compensates either for perturbations in the growth rate *or* the cell size, as a result of noisy cell division. Since our data also indicates a strong correlation between the growth rate of the cell and the protein synthesis rate, we speculate that this unidentified regulatory mechanism might control the growth rate by acting on the protein synthesis rate. Translation regulation in *B. subtilis*^29^ is not yet as well understood as in *E. coli*^30^, but its workings are qualitatively similar. In *E. coli* growth rate is adjusted by regulation of the ribosome concentration via ppGpp, a second messenger that is produced when translation is limited by amino acids^31^. This mechanism might provide a direct coupling between the metabolic state, perturbed by cell division, the balance between amino acid synthesis and consumption, during translation, and the cell’s growth rate. This coupling would lead to a strong correlation between the growth rate and the protein production rate of a cell, which is indeed what was observed (Fig. 4C). Since these two rates are strongly correlated, protein synthesis and dilution remain balanced, ensuring that the concentration of stable proteins remains homeostatic (Fig. 4D).

We observed deviations from a fixed exponential growth rate as cells progress through their cell cycle (Fig. 2B). Figure 3B indicates that this is because the absolute growth is not proportional to size. This becomes particularly evident when elongation rate is plotted versus time-to-division, allowing two distinct growth regimes or phases to be defined (Fig. 3C). Despite deviations from fixed exponential growth during the cell cycle, and significant size and growth rate differences at birth, all cells behave very similarly at the end of the cell cycle (Fig. 2B). By looking at growth dynamics during each of the identified phases, we could gain further insight into how growth rate and size compensations are achieved during the cell cycle.

In the first phase, during which the cells act as sizers (Fig. 5C), the elongation rate is constant and independent of the birth size of the cells. The fact that the specific growth rate varies with size (Fig. 2B) is therefore completely explained by the normalisation of this growth rate measure by length. The second phase has a fairly constant duration and the cells therefore behave as timers (Fig. 5D). Overall, they behave as imperfect adders (Fig. 5A) in agreement with previous findings^13^. Thus, the length added during phase I correlates strongly with size at birth (Fig. 5C). This correlation is much weaker for phase II, likely because small and large cells have converged in length when the second phase commences. As a consequence, while the average cell does add a fixed length per generation, smaller cells add more than average, and bigger cells less (Fig. 5A); this deviation from the pure adder-behaviour is similar to that reported by Wallden et al.^15^. An explanation for this size-added-per-generation bias could, as suggested for *E. coli,* lie in the coordination of cell mass with the initiation of DNA replication^15,17^; a size-threshold has to be reached before DNA replication can be initiated^15^. Consequently, smaller cells have to accumulate more mass before they reach this threshold. And so, if the duration of growth is approximately fixed after the initiation of DNA replication, the smaller a cell is at birth, the more total length it will have added by the time it divides again^15,17^.

A likely determinant of the growth rate of a cell is its protein synthesis rate, according to Fig. 4A-C. We note that Fig. 4C is in agreement with the growth-rate law of Maaløe^32^ and Hwa^33^, which shows a linear relation between the ribosomal protein fraction (proportional to the ribosome concentration) and the cell’s specific growth rate. Our results are in agreement with the growth law when the translation rate per ribosome is constant, i.e. when ribosomes operate at fixed saturation, as is expected from theory^34^.

Summarising, this work shows that cell division causes significant perturbations of the physiology of single *B. subtilis* cells, leading to large variations in the specific elongation rate and size of newborn cells. Perturbations are compensated for during the cell-cycle, and the cell-cycle can be decomposed into two phases: a first phase, of variable duration and a constant absolute elongation rate, during which cells compensate for size differences at birth, while the second phase lasts for a comparable duration across cells and all cells experience a very similar elongation rate increase. This suggests that growth rate and cell-size homeostasis are likely under a coordinated control mechanism that we do not yet mechanistically understand. Another finding of this work is that homeostasis of stable protein expression is achieved in *B. subtilis* by coordinated changes in cellular growth rate, which is responsible for protein concentration reduction via dilution, and protein synthesis. Single cells can differ significantly from each other immediately after birth, and this necessitates the activity of incompletely understood compensatory mechanisms that function to restore the behaviour of individual cells to that of the average cell. The metabolic and growth behaviour of single cells is therefore likely far removed from that of the average cell^35^. Most of our current knowledge of cellular metabolism still derives from population based methods and how it functions in single cells, in the context of cell-cycle progression, is an important next challenge.

## Methods

**Table 1:** Notations

| Notation | Description | Units |
| --- | --- | --- |
| $a$ | cell age, time elapsed since birth of a cell | h |
| $L$ | Length of a cell | $\mu\text{m}$ |
| $ER\vert a$ | Instantaneous elongation rate, $\frac{dL}{dt}$, of a cell at age $a$ | $\mu\text{m h}^{-1}$ |
| $\langle ER\vert a \rangle$ | Average ER, $\frac{dL}{dt}$, over all cells of age $a$ | $\mu\text{m h}^{-1}$ |
| $sER\vert a$ | Specific ER, $\frac{1}{L} \frac{dL}{dt}$, of a cell at age $a$ | $\text{h}^{-1}$ |
| $\langle sER\vert a \rangle$ | Average $sER$, $\frac{1}{L} \frac{dL}{dt}$, over all cells of age $a$ | $\text{h}^{-1}$ |
| $\overline{sER}$ | $sER$ averaged over the cell-cycle of a single cell | $\text{h}^{-1}$ |
| $\langle sER \rangle$ | mean $sER$ of the population | $\text{h}^{-1}$ |
| $T$ | generation time | h |
| $\alpha$ | Normalised cell age, $\frac{a}{T}$, of a cell | 0-1 |
| $L_b$ | Birth length of a cell | $\mu\text{m}$ |
| $L_d$ | Division length of a cell | $\mu\text{m}$ |
| $\langle x\vert \alpha \rangle$ | $x$ conditioned on $\alpha$ averaged over the population | $[x]$ |
| $\langle x\vert L_b, \alpha \rangle$ | $x$ subsetting by $L_b$, conditioned on $\alpha$ averaged over the $L_b$ subset | $[x]$ |
| Fluorescence (f) (per cell) | Sum of all pixel fluorescence intensities inside a cell | AU |
| Fluorescence concentration | Average pixel fluorescence intensity $\frac{f}{N_{pixels}}$ inside a cell | AU |

### Strains and medium composition

For growth experiments, prototrophic *Bacillus subtilis* strain BSB1^36^ was revived in a defined morphilinopropanesulphonic acid (MOPS) - buffered minimal medium (MM) containing: 40 mM MOPS (adjusted to pH 7.4), 2 mM potassium phosphate (pH 7.0), 15 mM (NH_4_)_2_SO_4_, and a trace element solution (final concentrations: 811 *μ*M MgSO_4_, 80 nM MnCl_2_, 5 *μ*M FeCl_3_, 10 nM ZnCl_2_, 30 nM CoCl_2_ and 10 nM CuSO_4_)^37^. Tris-Spizizen-salts (TSS) minimal medium composition was as following: 37.4 mM NH_4_Cl, 1.5 mM K_2_HPO_4_, 49.5 mM TRIS, 1mM MgSO_4_, 0.004% FeCl3 / 0.004% Na_3_-citrate*2H_2_O^38^ and trace elements as in the MM-medium. For solid TSS medium, 1.5% w/v low melt agarose was added.

The media were supplemented with different carbon sources to the following final concentrations: 6 mM arabinose, 5 mM glucose and 5 mM glucose with the amino acids methionine, histidine, glutamate and tryptophan to a final concentration of 1 mM each. We refer to these media as arabinose, glucose and glucose + 4 a.a., respectively. From a 1 M stock solution of isopropyl *β*-D-1-thiogalactopyranoside (IPTG) an appropriate amount was added to the medium to reach a final concentration of 50 or 1000 *μ*M.

*Escherichia coli* strain JM109 (Promega) was used for cloning and amplification of plasmids. For cloning, *E. coli* and *Bacillus subtilis* were grown in LB + 0.5% w/v glucose supplemented with the appropriate antibiotic in the following concentrations: ampicillin, 100 *μ*g/ml; spectinomycin 150 *μ*g/ml. For LB plates, 1.5% w/v agar were added prior to autoclaving.

Plasmid pDR111-N015-superfolderGFP was constructed by amplifying the coding sequence of superfolderGFP (sfGFP) by PCR with primers N015 (ggtggtgctagcaggaggtgatccagtatgtctaaaggtgaagaactg) and N017 (ggtggtgcatgcttatttgtagagctcatccat), digestion of the product and backbone pDR111^39^ (*bla amyE’ spc^R^ P_hyperspank_ lacI ’amyE*; kind gift from David Rudner) with NheI and SphI and subsequent ligation. After transformation of chemocompetent *Escherichia coli* JM109 (Promega) and plasmid isolation, the identity of pDR111-N015-sfGFP was confirmed by sequencing. *Bacillus subtilis* strain B15 (BSB1 *spc^R^ P_hyper–spank_^-^ sfGFP lacl::amyE*) was constructed as following: pDR111-N015-sfGFP was linearised with SacII, added to a BSB1 culture grown in MM+glucose until starvation phase, and incubated for one hour before addition of fresh MM and plating on LB+glucose+spc for selection. Genomic insertion into *amyE* was confirmed by amylase deficiency, PCR and sequencing. The *amyE* locus is situated at ≈ 28° on the genome.

### Growth experiments

Cells were revived by inoculation directly from single-use 15% glycerol stocks into 50 ml Greiner tubes with 5 ml MM supplemented with IPTG and grown at 37° Celsius and 200 rpm. After 8 to 15 generations, the cultures reached an OD_600_ between 0.01 and 0.2 and were diluted in 50 ml Greiner tubes with 5 ml liquid TSS supplemented with IPTG and grown at 37° Celsius and 200 rpm for another 4 to 5 generations. After dilution to an OD_600_ of 0.01, 2 *μ*l of the culture was transferred to a 1.5% low melt agarose pad freshly prepared with TSS.

Once seeded with cells, agarose pads were inverted and placed onto a glass bottom microwell dish (35 mm dish, 14 mm microwell, No. 1.5 coverglass) (Matek, USA), which was sealed with parafilm and immediately taken to the microscope for time-lapse imaging. Per carbon source, we carried out 5 independent growth experiments: *B. subtilis* B15 with 0, 50 and 1000 *μ*M IPTG and the parent strain *B. subtilis* BSB1 with 0 and 1000 *μ*M IPTG. For each experiment, we monitored growth at 4 different positions on the agarose pad. For the analysis of the growth dynamics, we combined the data from all 5 independent experiments per carbon source, as we did not detect significant differences in the specific elongation rates, length at birth and length at division between strains and conditions (Fig. S10). The analysis of protein expression was carried out on the data sets of *B. subtilis* B15 at full induction (1000 *μ*M IPTG).

### Microscopy and data analysis

Imaging was performed with a Nikon Ti-E inverted microscope (Nikon, Japan) equipped with 100X oil objective (Nikon, CFI Plan Apo λ NA 1.45 WD 0.13), Zyla 5.5 sCmos camera (Andor, UK), brightfield LED light source (CoolLED pE-100), fluorescence LED light source (Lumencor, SOLA light engine), GFP filter set (Nikon Epi-Fl Filter Cube GFP-B), computer controlled shutters, automated stage and incubation chamber for temperature control. Temperature was set to 37°C at least three hours prior to starting an experiment. Nikon NIS-Elements AR software was used to control the microscope. Brightfield images (80 ms exposure time at 3.2% power) were acquired every minute for 8-15 hours. GFP fluorescence images (1 second exposure at 25% power) were acquired every 10 min. Time-lapse data were processed with custom MATLAB functions developed within our group^4^. Briefly, an automated pipeline segmented every image, identifying individual cells and calculating their spatial features. Cells were assigned unique identifiers and were tracked in time, allowing for the calculation of time-dependent properties including cell ages, cell sizes (areas and lengths), elongation rates and generation times. In addition, the genealogy of every cell was recorded. The fluorescence values that we report here are the sum of all pixel intensities in the area of a cell contour. As a measure for fluorescence concentration we calculated the average pixel intensity in a fixed area in the centre of the cell. The output from the MATLAB pipeline was further analysed with MATHEMATICA, version 11 (Wolfram Research, Champaign, IL, USA), using custom scripts.

We excluded cells based on the following criteria: cells longer than 25*μ*m at division were excluded (filamentation), cells with a generation time longer than 250 minutes were excluded (sporulating cells), cells that only grew by 20% of their initial length (artifacts), cells that grew more than 4 times their initial length (filamentation) were excluded from the analysis. The cells removed in this way constituted less than 1% of each dataset.

#### Calculation of instantaneous rates

Instantaneous (specific) elongation rate was calculated using a sliding-window approach. Briefly, a window of a given size (5, 10 or 15 minutes for fast, intermediate and slow growth, respectively) was moved along the time series of length measurements for each cell and the difference or Log-difference of the first and last data point of the window was calculated as elongation rate or specific elongation rate, respectively. This resulted in a time-series of instantaneous (specific) elongation rates for each cell. The average of each of these time-series per cell is reported as 〈*ER*〉 or 〈*sER〉.*

Protein production rate was calculated similarly with a window size of 10 minutes, i.e. every time point where fluorescence was recorded, under all conditions. Protein capacity was obtained by normalising the protein production rate by cell length.

#### Binning by age

We binned time series data of all cells into 10 age bins of width 0.1 each. To avoid sampling bias resulting from varying interdivision times (IDT), i.e. overrepresentation of cells with long IDT per age bin due to fixed imaging intervals, we first binned the time series of individual cells into 10 age bins and averaged over each bin, before binning the data over the whole population/ birth length class.

#### Estimation of the growth phase transition prior to division

We estimated the time-to-division at which the elongation rate increases (Fig. 2e) by fitting a piecewise function to the data, using Wolfram MATHEMATICA’s *FindFit* function. The piecewise function that we used describes the elongation rate as function of time-to-division and assumes that elongation rate is constant initially (*a*) and increases exponentially at a certain time-to-division (*t_D_*):

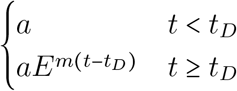

The function was fitted to the ensemble of data points from each birth length class, yielding 5 fits per growth condition.

### Correction for fluorescence drift over the time course of an experiment

To correct for an increase of background fluorescence that occurred over the time course of an experiment, we normalised the fluorescence values of each experiment by estimating the background fluorescence over time (for an example see Fig. S11). For this, per position, we defined 2 regions where no cell growth occurred and measured the evolution of fluorescence over the duration of the experiment. Using MATLAB, we fitted a polynomial to the rescaled background fluorescence and normalised the fluorescence values of all cells by the fitted function.

## Author contributions

NN, JvH and FB designed the study; NN performed experiments; JvH supplied custom software; NN and JvH performed analyses; NN, JvH and FB drafted and wrote manuscript.

## Appendices

### Theory

#### Elongation rate of a single cell as function of its cell-cycle

For single cells to grow balanced, their intracellular state, e.g. of metabolism, must be such that the cell’s specific growth rate 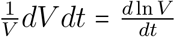 equals a constant denoted by *μ_V_*. The condition for this state is that all concentrations in a cell remain constant; since,

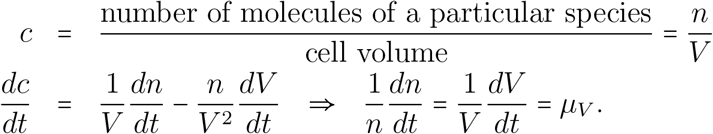

Thus, when the specific rate of molecule synthesis and volume are equal then concentrations are constant and will remain so and define the specific growth rate of a cell. Since, 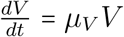 we obtain 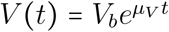. With *V_b_* as the birth volume of a cell and the generation time, *t_g_*, is defined as 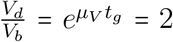, with *V_d_* as the division volume, such that 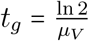.

The volume of an idealised rod-shaped cell equals the volume *V_cap_* of a sphere, with radius *r,* plus that of a cylinder with length *L* = l − 2*r*, with *l* as the length of the cell and *r* as its radius,

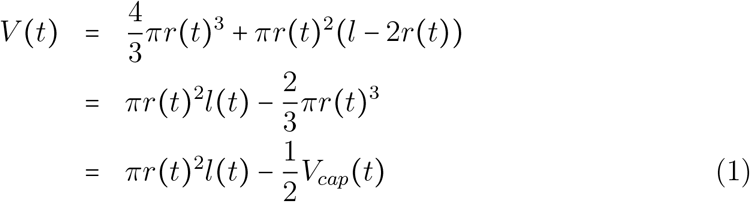

When we assume that the cell’s volume grows by length then *r*(*t*) becomes a constant *r* and

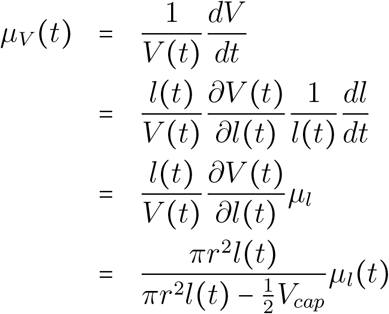

Thus when *μ_V_* is constant during balanced growth *μ_l_* is not; since,

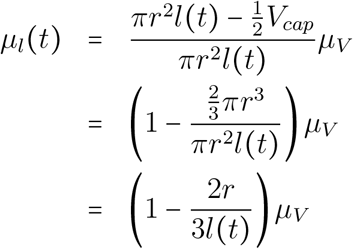

From equation 1 we can obtain *l*(*t*) in terms of *V*(*t*), for *V*(*t*) we can substitute *V*(*t*) = *V_b_e^μ_V_t^* and 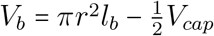, which gives for *l*(*t*) and *μ_l_*(*t*),

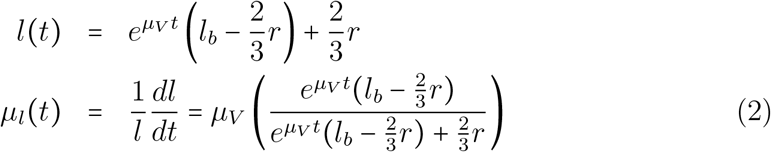

Note that *l_b_* ≥ 2*r* such that 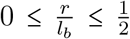 and that 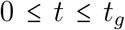. This last equation indicates that *μ_l_* ≠ *μ_V_* under conditions of balanced growth when *μ_V_* is fixed.

### Robustness of growth patterns to noise

To test whether the growth patterns and structured deviations form exponential growth we observe in our data is robust to measurement noise, we performed growth simulations by random sampling of measured growth parameters. For these simulations we assumed that single cells *<u>do</u>* grow exponentially during cell cycle progression, and that size homeostasis is achieved by a pure adder mechanism. First, we will outline the sampling and simulation procedure. Following this, we will compare simulated to experimentally measured growth data, using different single-cell profile alignment perspectives. This comparison serves demonstrates that the growth patterns we observe are not artifacts of different cell-cycle alignment perspectives when working with noisy single cell data.

#### Sampling and simulations

Random combinations of independently sampled (from experimentally measured distributions) growth parameters were combined and single cell growth trajectories were then simulated. Sampling was done without replacement. The data shown here is for the intermediate growth rate condition (glucose). The following variables were sampled (for glucose simulations 12553 unique combination were generated, the same number experimentally observed cells):

- Birth length (*μ*m)
- Added length (*μ*m, Division length - Birth Length)
- Average specific elongation rate, Mu (min^−1^), calculated from as the slope of a linear fit to the ln-transformed length profiles of cells

Using these sampled variables, individual growth trajectories, according to a pure-adder model, were simulated as follows:

1. Calculate (Division length)_*i*_ for the combination of randomly sampled (Birth length)_*i*_ and randomly sampled (Added length)_*i*_

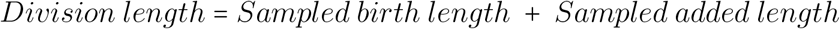
2. Calculate (generation time)_*i*_ (minutes) from the sampled (birth length)_*i*_, the calculated Division length (see 1) and the sampled (Mu)_*i*_ (min^−1^)

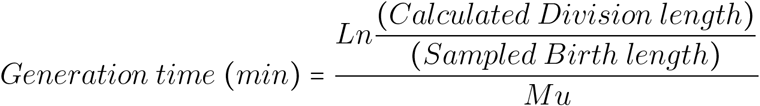
3. Simulate time-dependent length profiles (*L*(*t*)) for each of the 12553 randomly generated growth parameter set

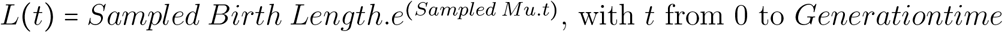

##### To summarise

The simulated cell trajectories shown below are generated from the glucose dataset. This dataset consists of 12553 individual cell trajectories. An equal number of simulated cell trajectories were generated from randomly drawn (with replacement) measured birth lengths, added lengths (i.e. division length birth length) and Mu (i.e. measurements are just randomly recombined). Histogram comparisons of measured vs simulated growth parameters are is shown in Fig. S1.

**Figure S1:**
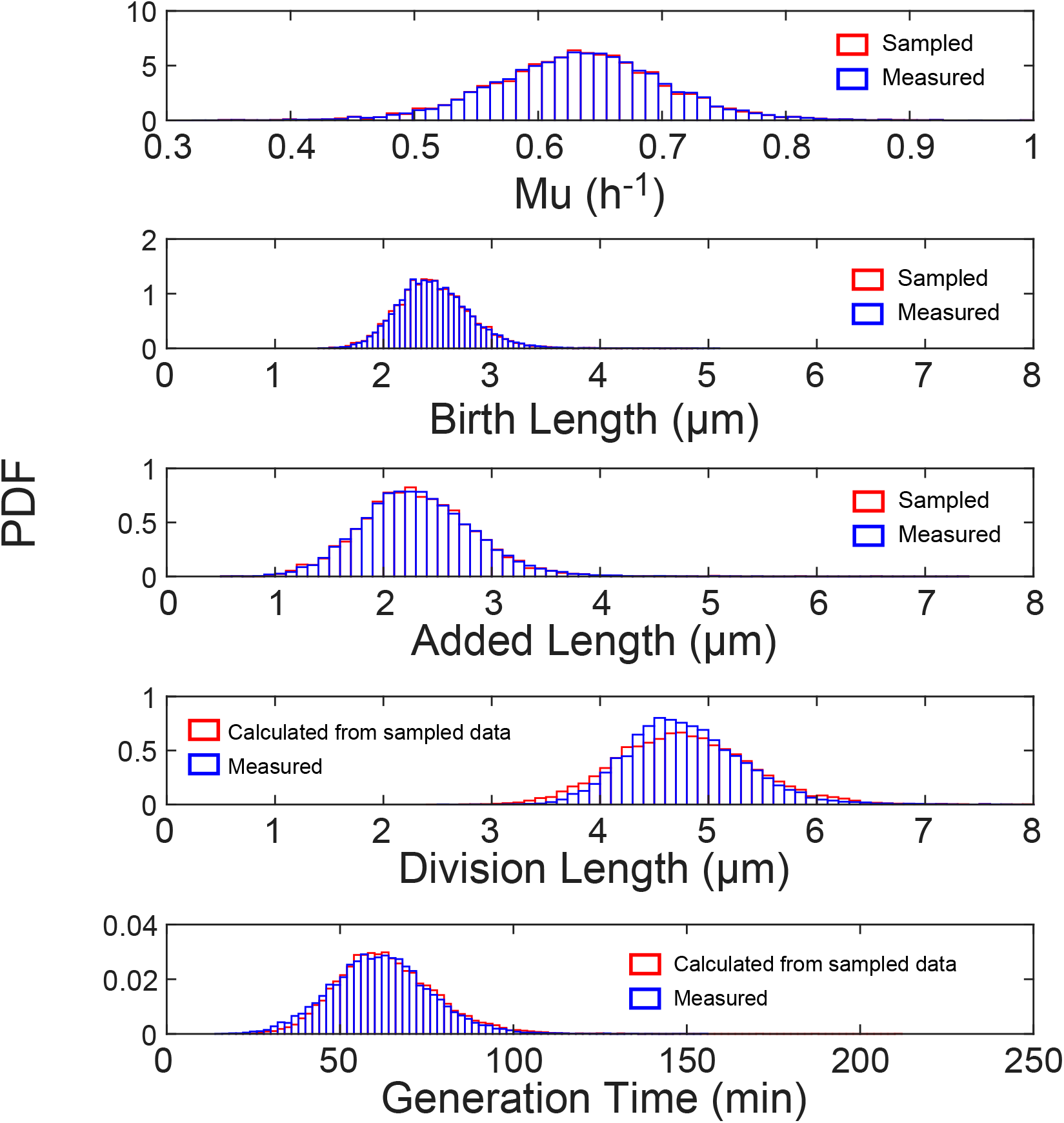
Comparison of measured vs simulated data. Distributions of sampled values (top three panels): Mu (Average specific elongation rate), Birth Length and Added Length. Simulated variables, Division length and Generation time, are shown in the bottom two panels. Mu is calculated as the slope of a linear fit to the ln-transformed length profile of a cell

#### The average cell and different length-profile alignment perspectives

##### Normalized/Rescaled Cell Cycle age

By rescaling (normalising) the generation times of individual cells (birth = 0 and division =1), the length profiles of cells with varying generation times can easily be aligned. However, this comes with the caveat that any dynamics (if present) will be compressed, for cells with longer than average generation times, or stretched for cells with shorter than average generation times. However, this rescaling-approach provides is the most commonly used method to directly compare the growth profiles of individual cells with varying generation times. Furthermore, it allows for the average-cell cycle profile to be studied and quantified. We find that the average cell-cycle profile shows structured deviations from exponential growth (Figure 1, main text). By quantifying the residuals of a linear fit to the ln-transformed average length profile of all cells, we a clear and structured deviations from exponential growth (Fig. S2). For comparison, these structured deviations are not present in the (noisy) simulated data (see above).

Next, we compare the experimentally determined specific elongation rate profile of the average cell, as function of rescaled age, to that of the simulated data (Fig. S3). The profiles essentially reflect the structure of the residuals shown in Fig. S2, with experimental data showing clear and structured cell-cycle-dependent deviations from exponential growth, while the simulated data does not. From these analyses, it is clear that the average cell-cycle growth pattern we observe it is not a consequence of the alignment (through rescaling of age) of noisy single cell data.

**Figure S2:**
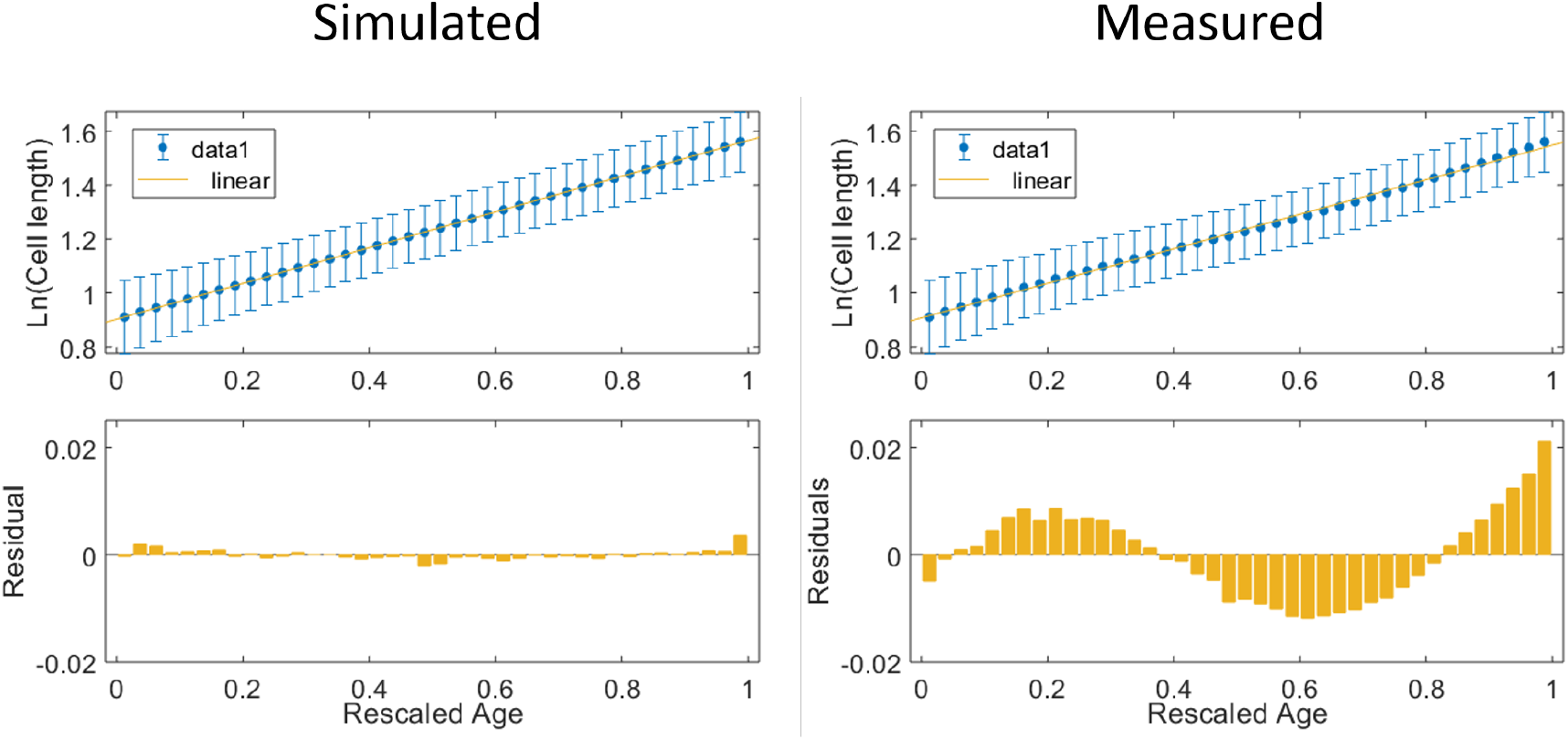
Average cell-length profiles. The Average ln-transformed cell length-profile is shown for 12553 simulated cells (top left panel) and 12553 measured cells (top right panel). Bottom panel shows the residuals of a linear-fit (yellow lines in top panels) to the ln-transformed cell length data. Cell length data was binned (bin size = 0.025) separately for each cell, using rescaled age, and then averaged across cells. Error bars show standard deviation of cell lengths at each age bin. Note that we use Rescaled age bin sizes of 0.1 for our main analysis, ensuring larger cell numbers per bin. Here a smaller bin size is used to imbue the residual plots with more data points and enhance the structured profile.

**Figure S3:**
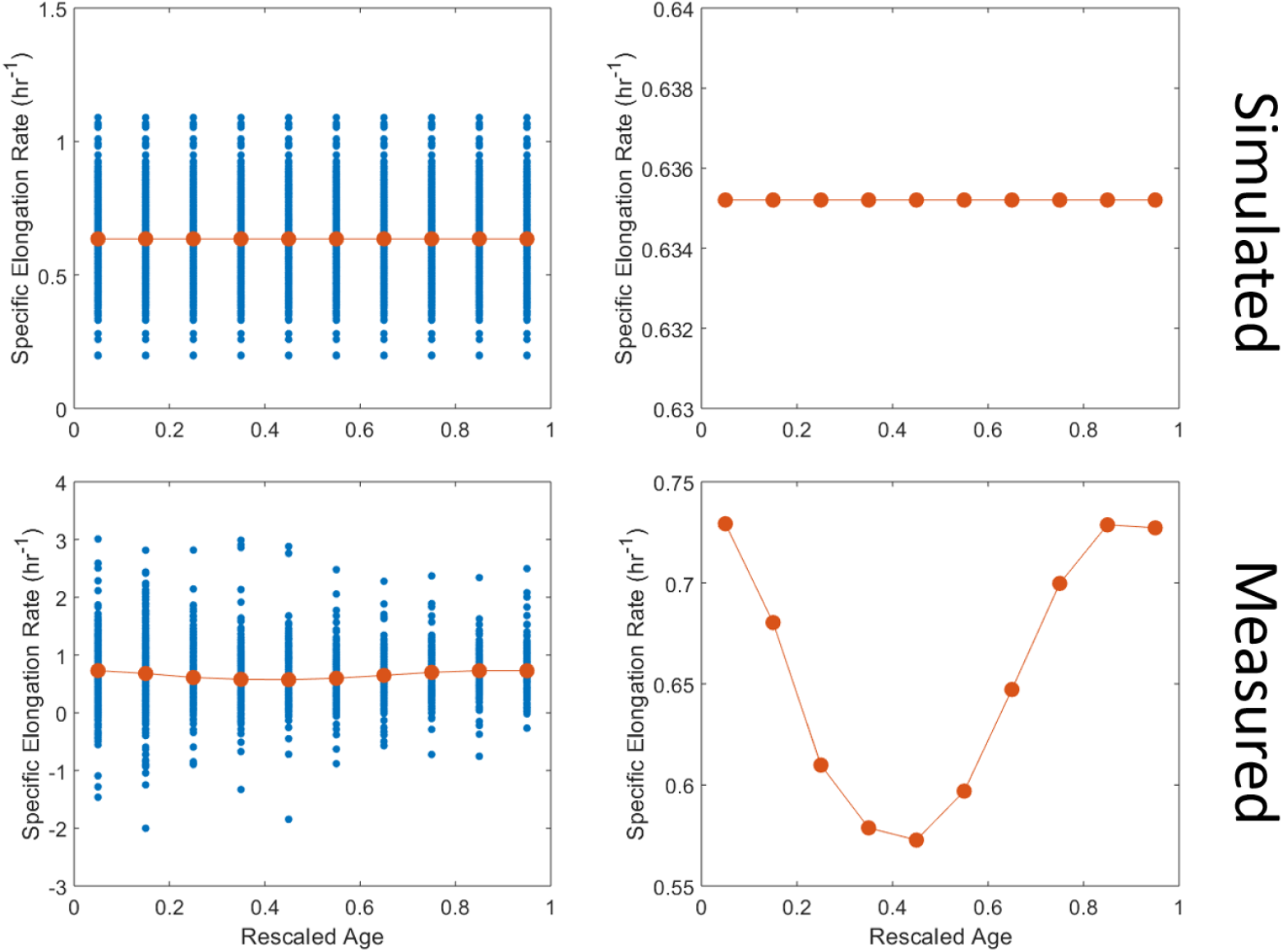
Average (piecewise) specific elongation rate profiles. Specific elongation rate is shown as function of Rescaled Age. Blue shows single cell data, red shows population average. As before, data is show for 12553 simulated cells (top panels) and 12553 measured cells (bottom panels). Rate data was binned (bin size = 0.1) separately for each cell, using rescaled age, and then averaged across cells.

##### Absolute age: time-since-birth vs time-to-division

We state in the main text, that by using time-to-division to anchor single cell profiles, a biphasic growth pattern becomes visible. Here we again compare (noisy) > simulated data with experimentally measured data (Fig. S4 and S5 and), and show that the patterns we observe are not a consequence of measurement noise being imbued with structure due to a specific alignment choice. What is clear is that the biphasic pattern is only visible when aligning cells according to their time-to-division, and then only in the experimental data. In Fig. S5, we overlay the time-to-division average elongation rate profiles of simulated and measured data, also indicating the phase transition point calculated for the glucose condition. It can clearly be seen that the simulated data does not display the biphasic pattern we observe experimentally.

**Figure S4:**
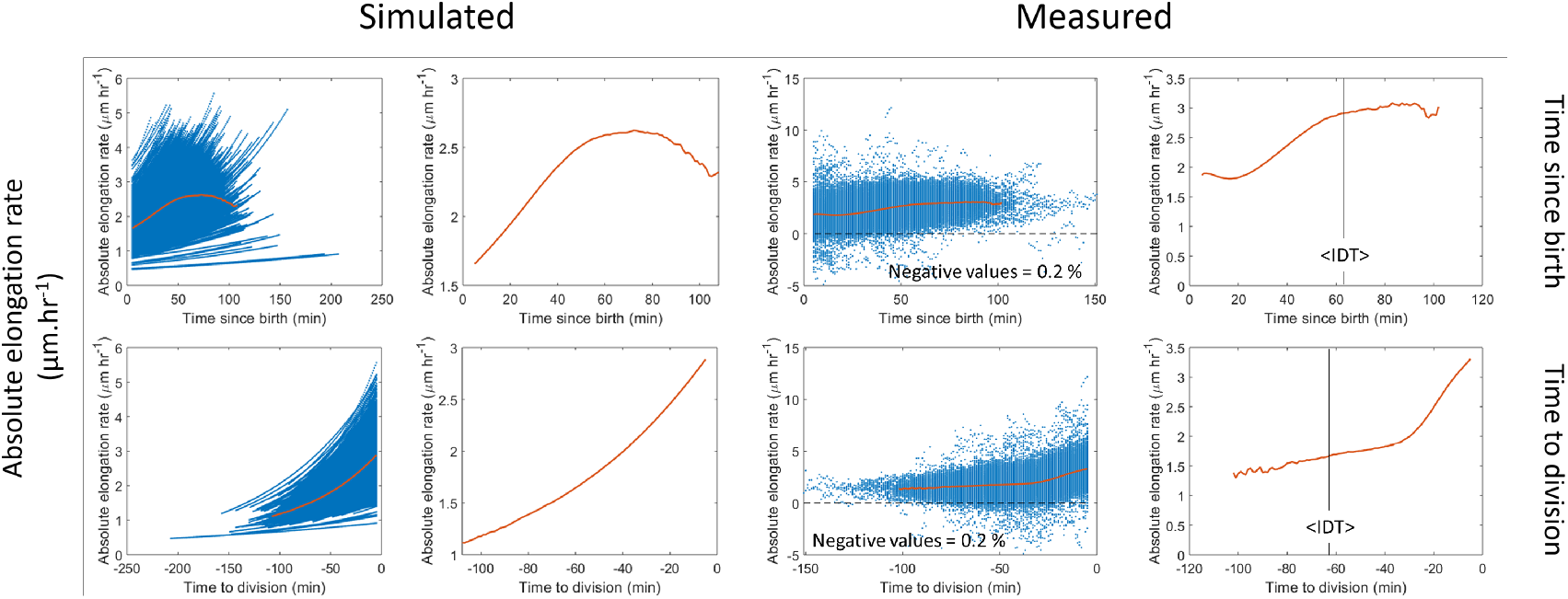
Comparison of Simulated vs Measured Absolute elongation rate profiles. For both Simulated and Experiment data, the Absolute elongation rates as function of either Time since Birth (Top panels) or Time to Division (bottom panels) are shown. In the experimental data some negative elongation rates are observed (i.e. cell shrinkage), but these are likely due to segmentation errors and constitute only 0.2 % of all measurements. Individual data points are indicated in blue and represent measurements form 12553 unique cells. The population averages are indicated by the orange lines and are shown both superimposed on single cell data, and in by separate panel (to the right of the single data) with a different y-axis scale to clearly show the average pattern. < *IDT* > indicates the average measured interdivision time (NOTE: Interdivision time is the same as the Generation time). Average profiles are calculated only for time points with at least 50 cells.

**Figure S5:**
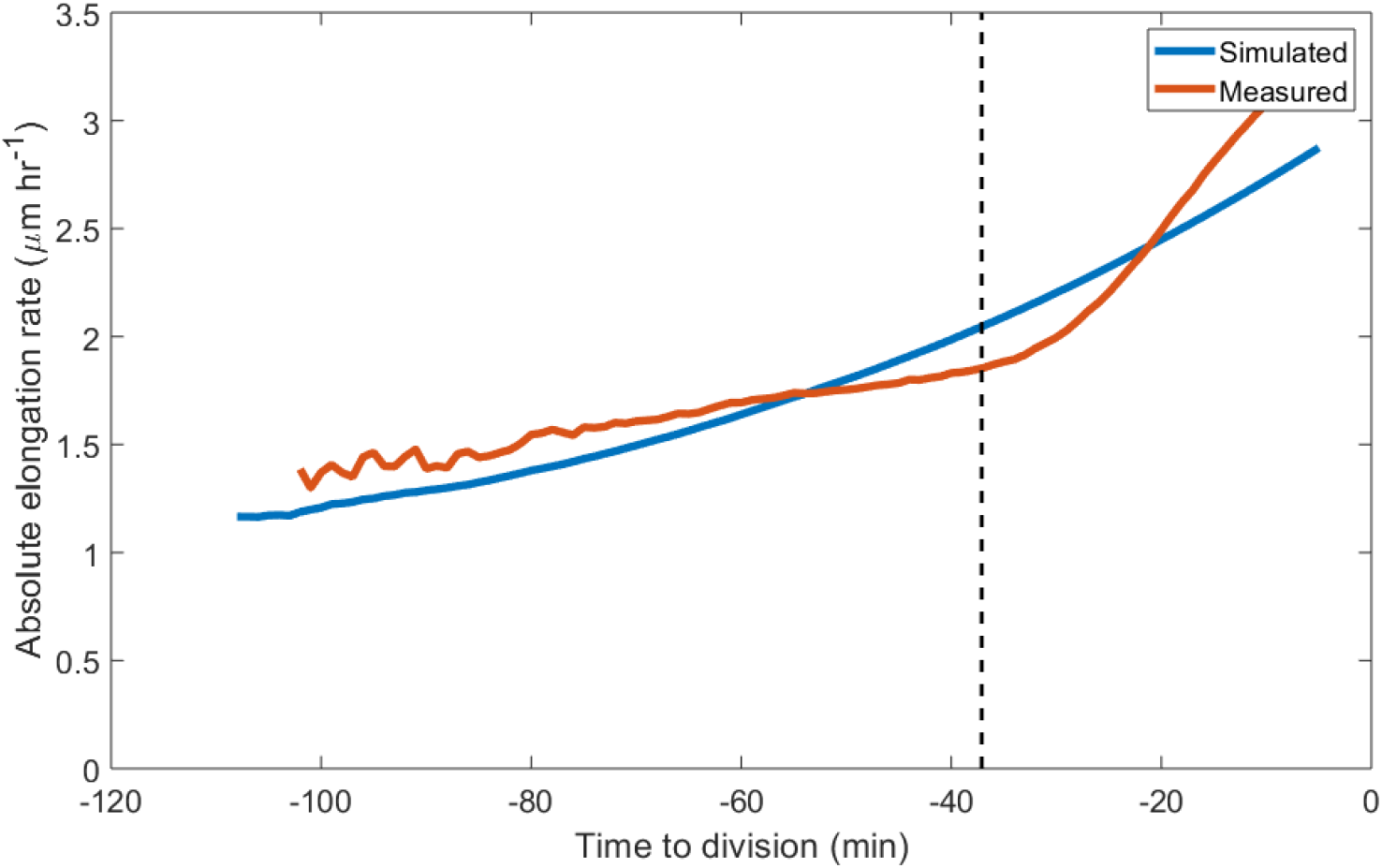
Direct comparison of population average simulated vs measured absolute elongation rate profiles. Data is shown for time points with at least 50 cells, and is the sames in Fig. S4. The dashed line shows the phase transition point as calculated from experimental data (see Materials and Methods in main manuscript). Average profiles are calculated only for time points with at least 50 cells.

**Figure S6:**
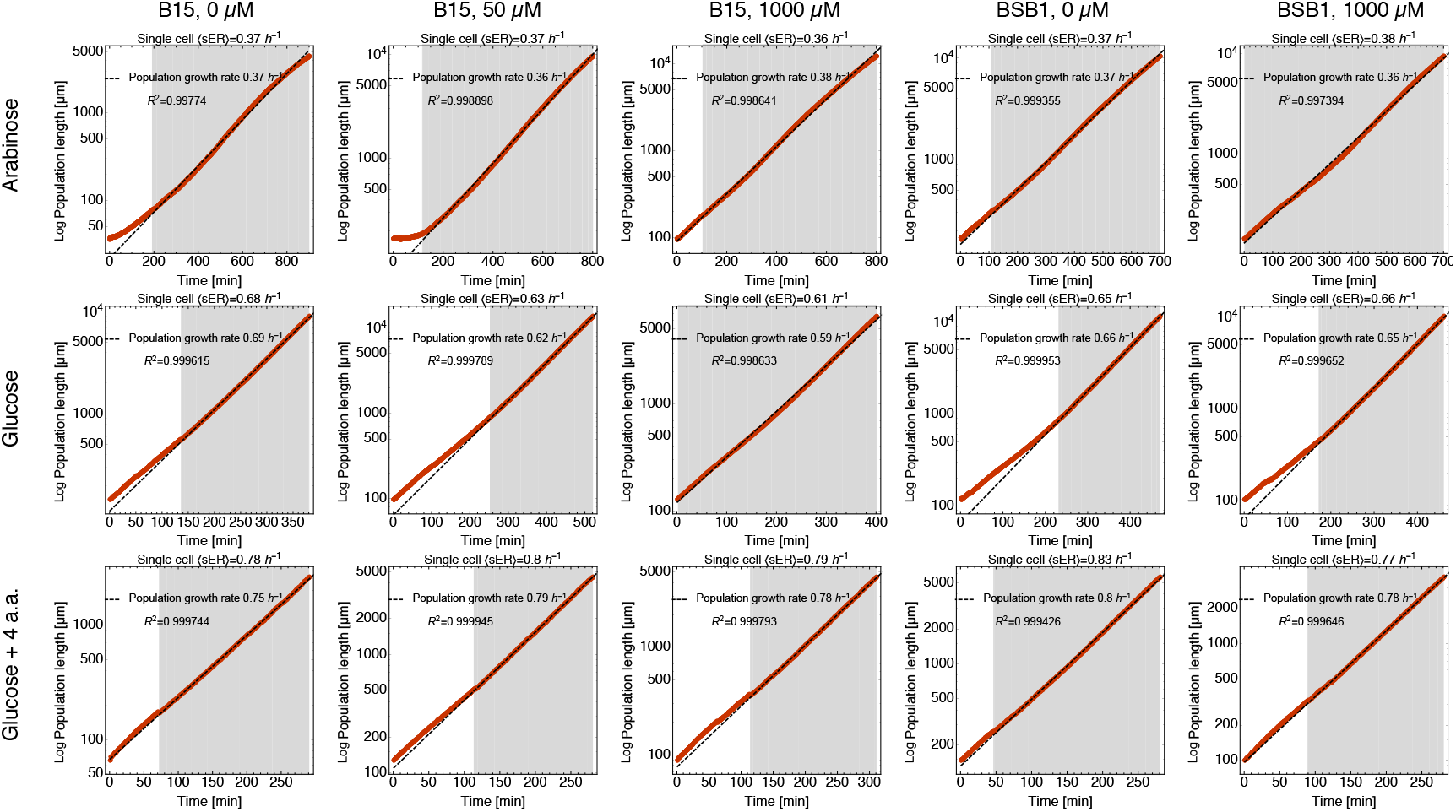
Comparison between single cell average 〈*sER*〉 and population growth rate as measured by the length increase of the whole population. The dashed line is a linear fit to the *log*-transformed total cell length of the population, the slope of which yields the specific growth rate. We define the region of exponential growth by an *R*^2^-cutoff (*R*^2^ ≥ 0.995) of the fit to the *log*-transformed data (grey area). In all analyses in the main text, we only consider cells that were born within the grey area.

**Figure S7:**
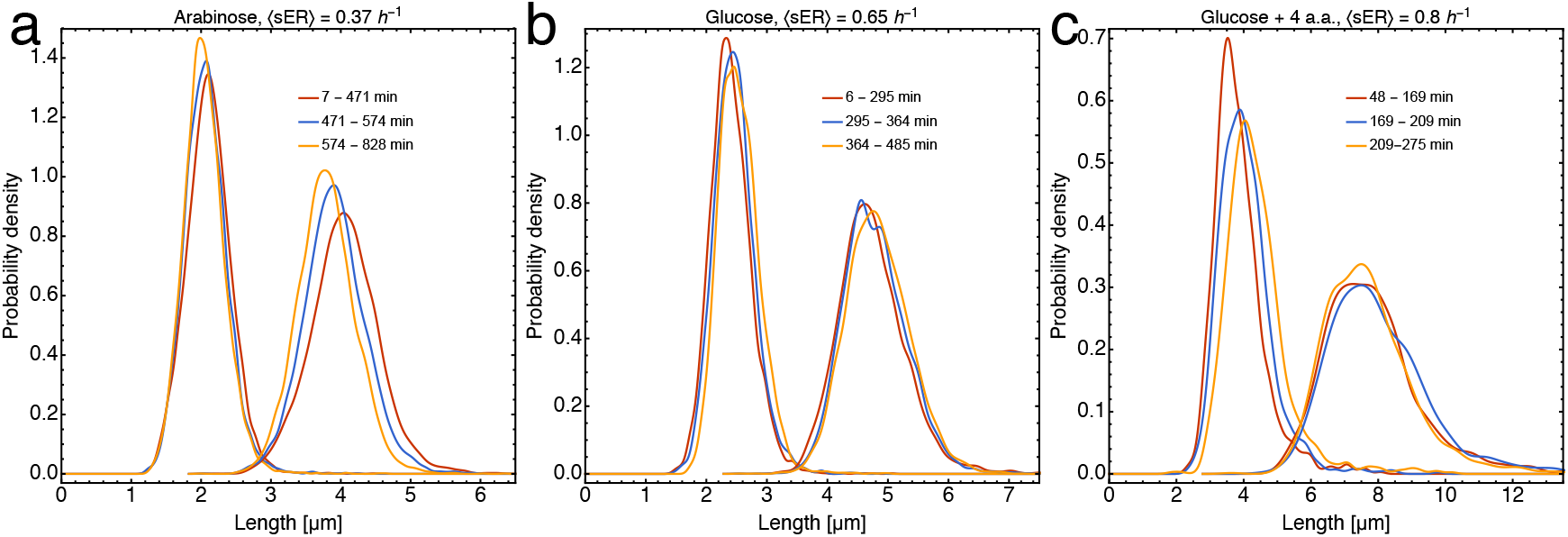
Distributions of birth length and division length are stable for several generations, indicating balanced population growth. Each distribution represents a third of all cells per condition from the period of balanced growth (cf. Fig. S6). The legend indicates the time frame in which cells from the respective distribution were born during the individual experiments.

**Figure S8:**
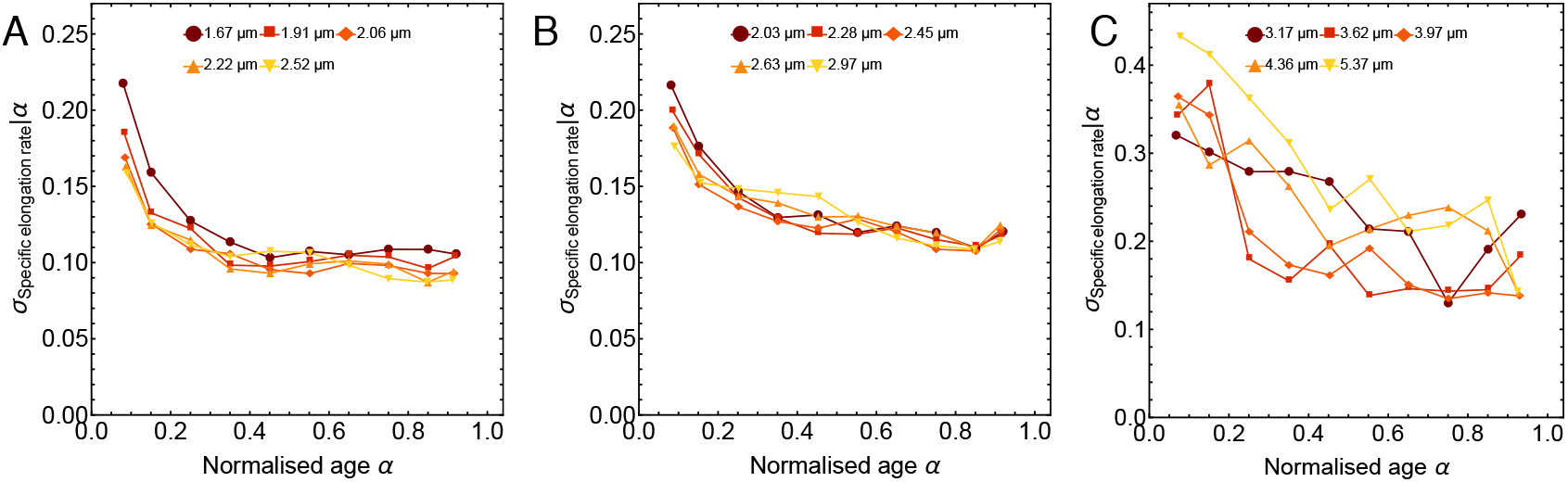
Variation in growth rate decreases as function of the cell-cycle. **A.** Standard deviation of the specific elongation rate for slow (0.37 *h*^−1^), **B.** intermediate (0.65 *h*^−1^) and **C.** fast (0.8 *h*^−1^) growth condition.

**Figure S9:**
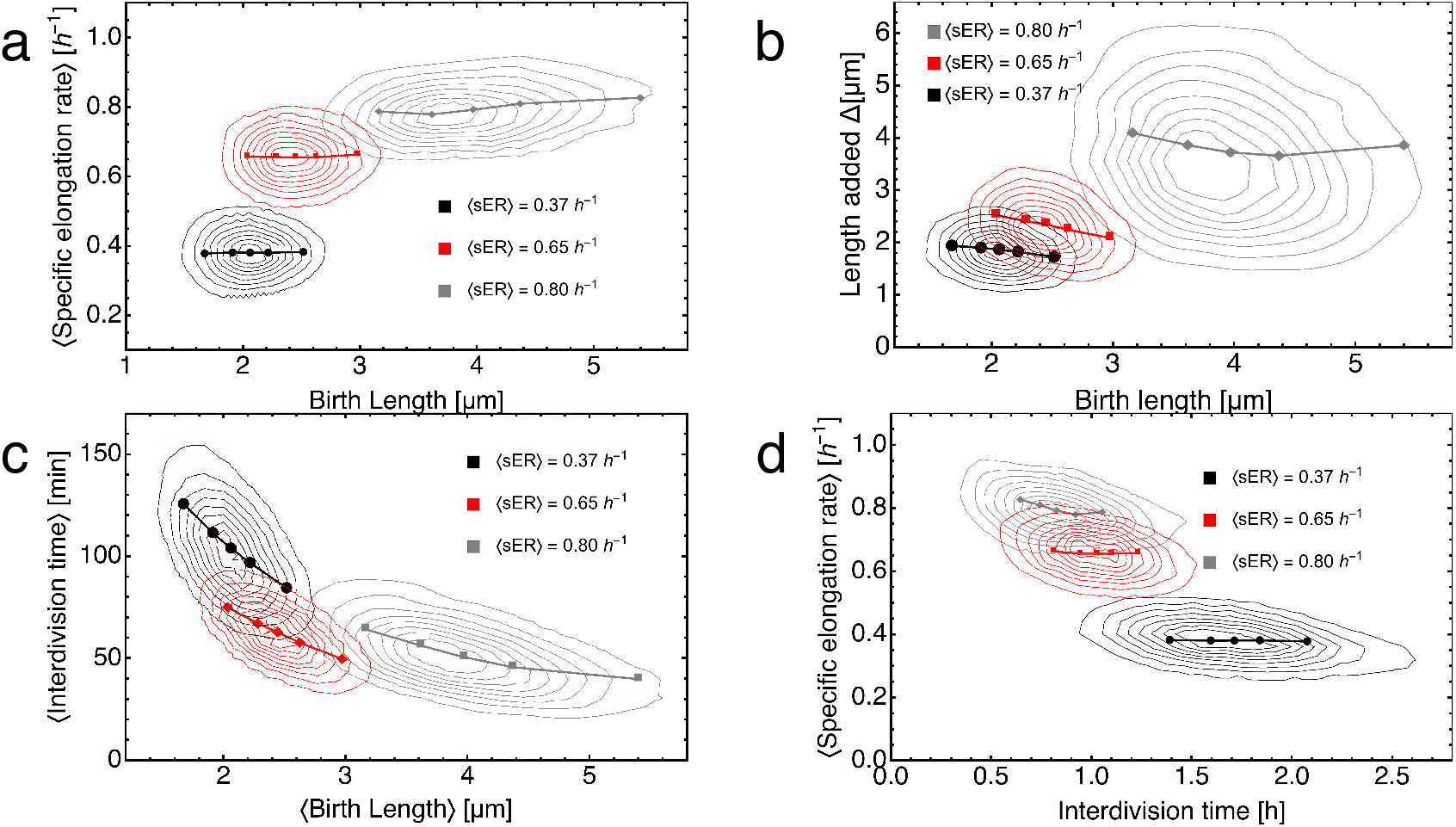
Correlations related to cell size homoeostasis. **a.** Specific elongation rate is independent of birth length. **b.** The amount a cell grows before division is negatively correlated with birth length, i.e. smaller cell grow more. **c.** Interdivision time decreases with increasing birth length. **d.** Interdivision time and specific elongation rate are hardly correlated. The symbols represent the different birth length classes from the main text.

**Figure S10:**
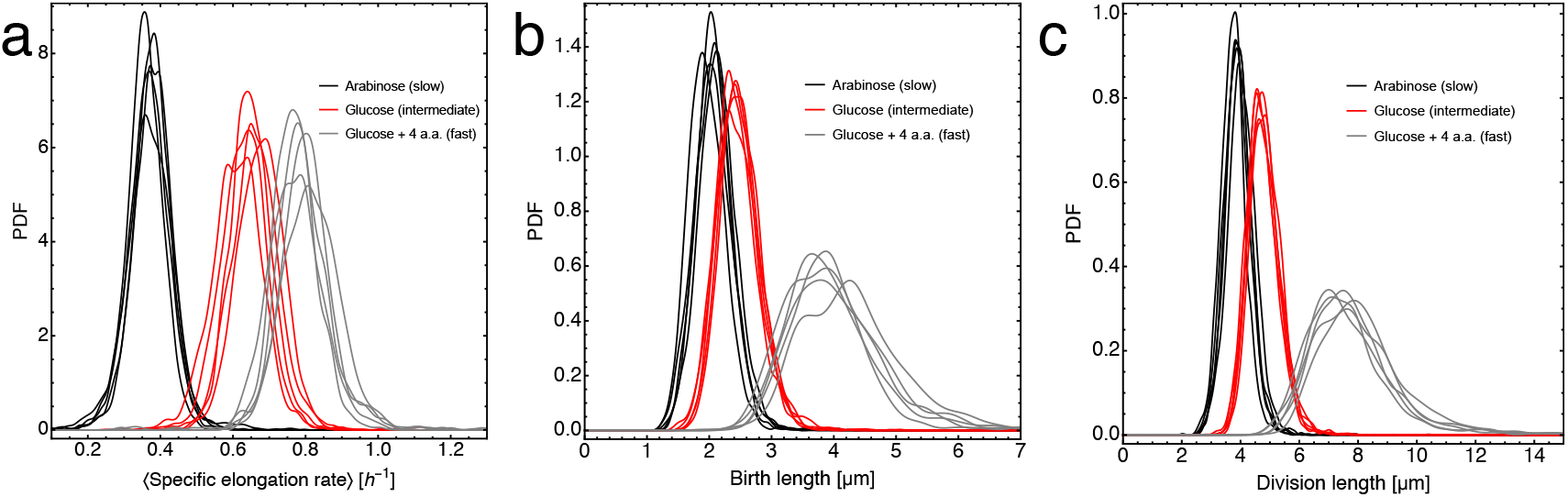
**A.** Distributions of specific elongation rate, **B.** birth lengths and **C.** division lengths for all 5 experiments per carbon source. All 5 experiments per carbon source were combined for the growth analyses the main text.

**Figure S11:**
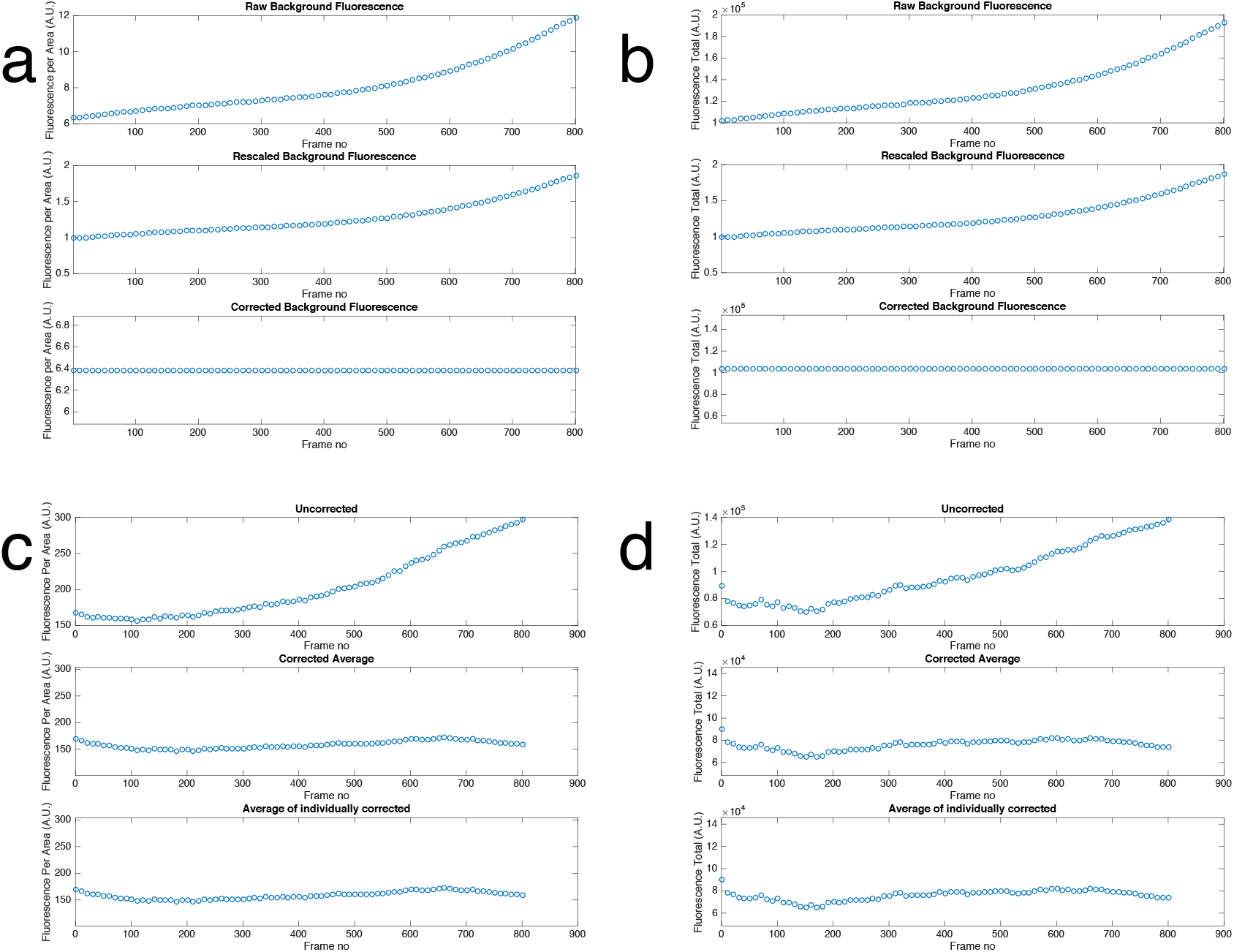
Correction for fluorescence drift. The drift in fluorescence over the course of each experiment was estimated from an area in the field of view without cell growth (**a., b.**, upper panel, see Methods for details). A polynomial function was fitted to the rescaled background fluorescence and used to normalise the fluorescence values of all cells (**c., d.**). The example shown here is for B15, grown on arabinose with 1000 *μ*M IPTG added.

